# The 100 € lab: A 3-D printable open source platform for fluorescence microscopy, optogenetics and accurate temperature control during behaviour of zebrafish, *Drosophila* and *C. elegans*

**DOI:** 10.1101/122812

**Authors:** Andre Maia Chagas, Lucia Prieto Godino, Aristides B. Arrenberg, Tom Baden

## SUMMARY

Small, genetically tractable species such as larval zebrafish, *Drosophila* or *C. elegans* have become key model organisms in modern neuroscience. In addition to their low maintenance costs and easy sharing of strains across labs, one key appeal is the possibility to monitor single or groups of animals in a behavioural arena while controlling the activity of select neurons using optogenetic or thermogenetic tools. However, the purchase of a commercial solution for these types of experiments, including an appropriate camera system as well as a controlled behavioural arena can be costly. Here, we present a low-cost and modular open-source alternative called “FlyPi”. Our design is based on a 3-D printed mainframe, a Raspberry Pi computer and high-definition camera system as well as Arduino-based optical and thermal control circuits. Depending on the configuration, FlyPi can be assembled for well under 100 € and features optional modules for LED-based fluorescence microscopy and optogenetic stimulation as well as a Peltier-based temperature stimulator for thermogenetics. The complete version with all modules costs ∼200 €, or substantially less if the user is prepared to “shop around”. All functions of FlyPi can be controlled through a custom-written graphical user interface. To demonstrate FlyPi’s capabilities we present its use in a series of state-of-the-art neurogenetics experiments. In addition, we demonstrate FlyPi’s utility as a medical diagnostic tool as well as a teaching aid at Neurogenetics courses held at several African universities. Taken together, the low cost and modular nature as well as fully open design of FlyPi make it a highly versatile tool in a range of applications, including the classroom, diagnostic centres and research labs.

## INTRO

The advent of protein engineering has brought about a plethora of genetically encoded actuators and sensors that have revolutionised neuroscience as we knew it but a mere decade ago. On the back of an ever-expanding array of genetically accessible model organisms, these molecular tools have allowed researchers to both monitor and manipulate neuronal processes at unprecedented breadth (e.g.: [1]–[3]). In parallel, developments in consumer-oriented manufacturing techniques such as 3-D printing as well as low-cost and user-friendly microelectronic circuits have brought about a silent revolution in the way that individual researchers may customise their lab-equipment or build entire setups from scratch (reviewed in: [4]–[7]). Similarly, already ultra-low cost light emitting diodes (LEDs), when collimated, now provide sufficient power to photo-activate most iterations of Channelrhodopsins or excite fluorescent proteins for optical imaging, while a small Peltier-element suffices to thermo-activate heat-sensitive proteins [8], [9]. In tandem, falling prices of high-performance charge-coupled device (CCD) chips and optical components such as lenses and spectral filters mean that today already a basic webcam, in combination with coloured transparent plastic or a diffraction grating, may suffice to perform sophisticated optical measurements [10], [11]. Taken together, modern biosciences today stand at a precipice of technological possibilities, where a functional neuroscience laboratory set-up capable of delivering high-quality data over a wide range of experimental scenarios can be built from scratch for a mere fraction of the cost traditionally required to purchase any one of its individual components. Here, we present such as design.

Assembled from readily available off-the-shelf mechanical, optical and electronic components, the “FlyPi” provides a modular solution for basic light- and fluorescence-microscopy as well as time-precise opto- and thermogenetic stimulation during behavioural monitoring of small, genetically tractable model species such as zebrafish (*Danio rerio*), fruit flies (*Drosophila Melanogaster*) or nematodes (e.g. *Caenorhabditis elegans*). The system is based on an Arduino microcontroller [12] and a Raspberry Pi 3 single board computer (RPi3; [13]), which also provides sufficient computing power for basic data analysis, word processing and web-access using a range of fully open-source software solutions that are pre-installed on the secure digital (SD)-card image provided. The mechanical chassis is 3-D printed and all source code is open, such that the design and future modifications can be readily distributed electronically to enable rapid sharing across research labs and institutes of science education. This not only facilitates reproducibility of experimental results across labs, but promotes rapid iteration and prototyping of novel modifications to adapt the basic design for a wide range of specialised applications. More generally, it presents a key step towards a true democratisation of scientific research and education that is largely independent of financial backing [4].

Here, we first present the basic mode of operation including options for micropositioning of samples and electrodes and demonstrate FlyPi’s suitability for light microscopy and use as a basic medical diagnostic tool. Second, we present its fluorescence capability including basic calcium imaging using GCaMP5 [1]. Third, we survey FlyPi’s suitability for behavioural tracking of *Drosophila* and *C. elegans.* Fourth, we demonstrate optogenetic activation of Channelrhodopsin 2 [3] and CsChrimson [14] in transgenic larval zebrafish as well as *Drosophila* larvae and adults. Fifth, we evaluate performance of FlyPi’s Peltier-thermistor control loop for thermogenetics [15]. Sixth, we briefly summarise our efforts to introduce this tool for university research and teaching in sub-Saharan Africa [4], [16].

## RESULTS

### Overview

The basic FlyPi can resolve samples down to ∼10 microns, acquire video at up to 90 Hz and acquire time-lapse series over many hours. It consists of the 3-D printed mainframe (Fig. 1A-D), one RPi3 computer with a Pi-camera and off-the-shelf objective lens, one Arduino-Nano microcontroller as well as a custom printed circuit board (PCB) for flexible attachment of a wide range of actuators and sensors (Fig. 1C). The main printed frame allows modular placement of additional components into the camera-path such as holders for petri-dishes (Fig. 1H) or microscope slides (Fig. 1I). This basic build, including power adapters and cables and the module for lighting and optogenetic stimulation can be assembled for <100 € (Supplementary Table 1; Fig. 1D). Additional modules for fluorescence imaging (Fig. 1E, cf. Fig. 3), temperature control (Fig. 1F, cf. Fig. 6) or an automated focus drive (Fig. 1G) can be added as required. For a full bill of materials (BOM), see Supplementary Table 1. A complete user manual and assembly instructions are included in the Supplementary materials.

**Figure 1.**
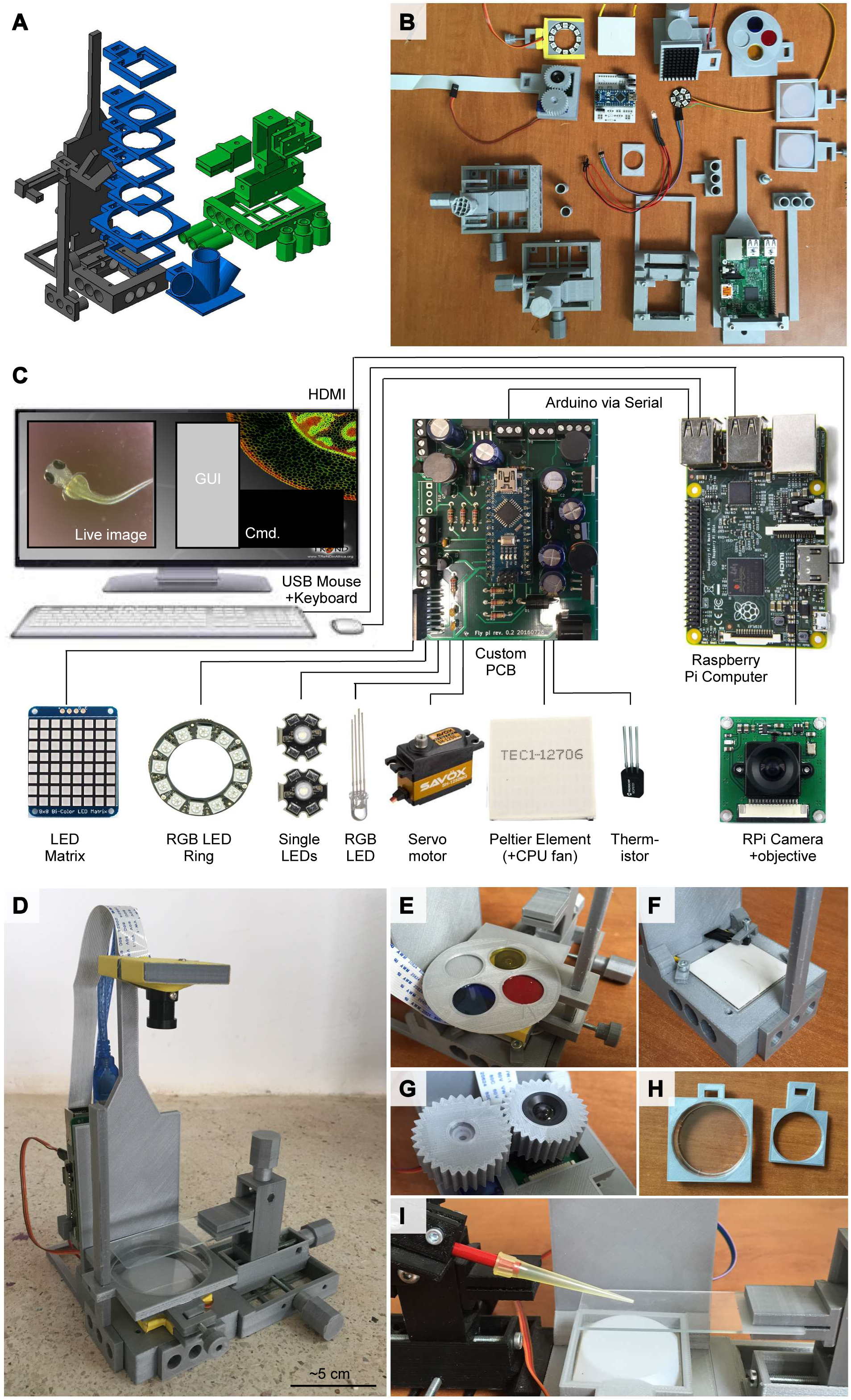
Overview. **A,** The 3D model, colour coded by core structure (black), mounting adapters (blue) and micromanipulator (green). **B**, Printed parts and electronics, part-assembled. **C**, Wiring diagram and summary of electronics. Full bill of materials (BOM) in Supplementary Table 1. **D**, The assembled FlyPi with single micromanipulator and LED-ring module, diffusor and Petri-dish adapter mounted in the bottom. **E**, Filter wheel mounted above the inverted camera objective. **F**, Peltier element and thermistor embedded into the base. **G**, Automatic focus drive. **H**, Petri-dish mounting adapters. **I**, A second micromanipulator mounted to the left face of FlyPi holding a probe (here, a 200 μl pipette tip for illustration) above the microscope slide mounted by the micromanipulator on the right.

### Basic camera operation and microscopy

To keep the FlyPi design compact and affordable yet versatile, we made use of the RPi platform, which offers a range of FlyPi-compatible camera modules. Here, we use the “adjustable focus RPi RGB camera” (Supplementary Table 1) which includes a powerful 12 mm threaded objective lens. Objective focal distance can be gradually adjusted between ∼1 mm (peak zoom, cf. Fig. 2D) and infinity (panoramic, not shown), while the camera delivers 5 megapixel Bayer-filtered colour images at 15 Hz. Spatial binning increases peak framerates to 42 Hz (x2) or 90 Hz (x4). Alternatively, the slightly more expensive 8 megapixel RPi camera or the infrared-capable NO-IR camera can be used. Objective focus can be set manually, or via a software-controlled continuous-rotation micro servo motor (Fig. 1G). Alternatively, the RPi CCD chip can be directly fitted above any other objective with minimal mechanical adjustments.

**Figure 2.**
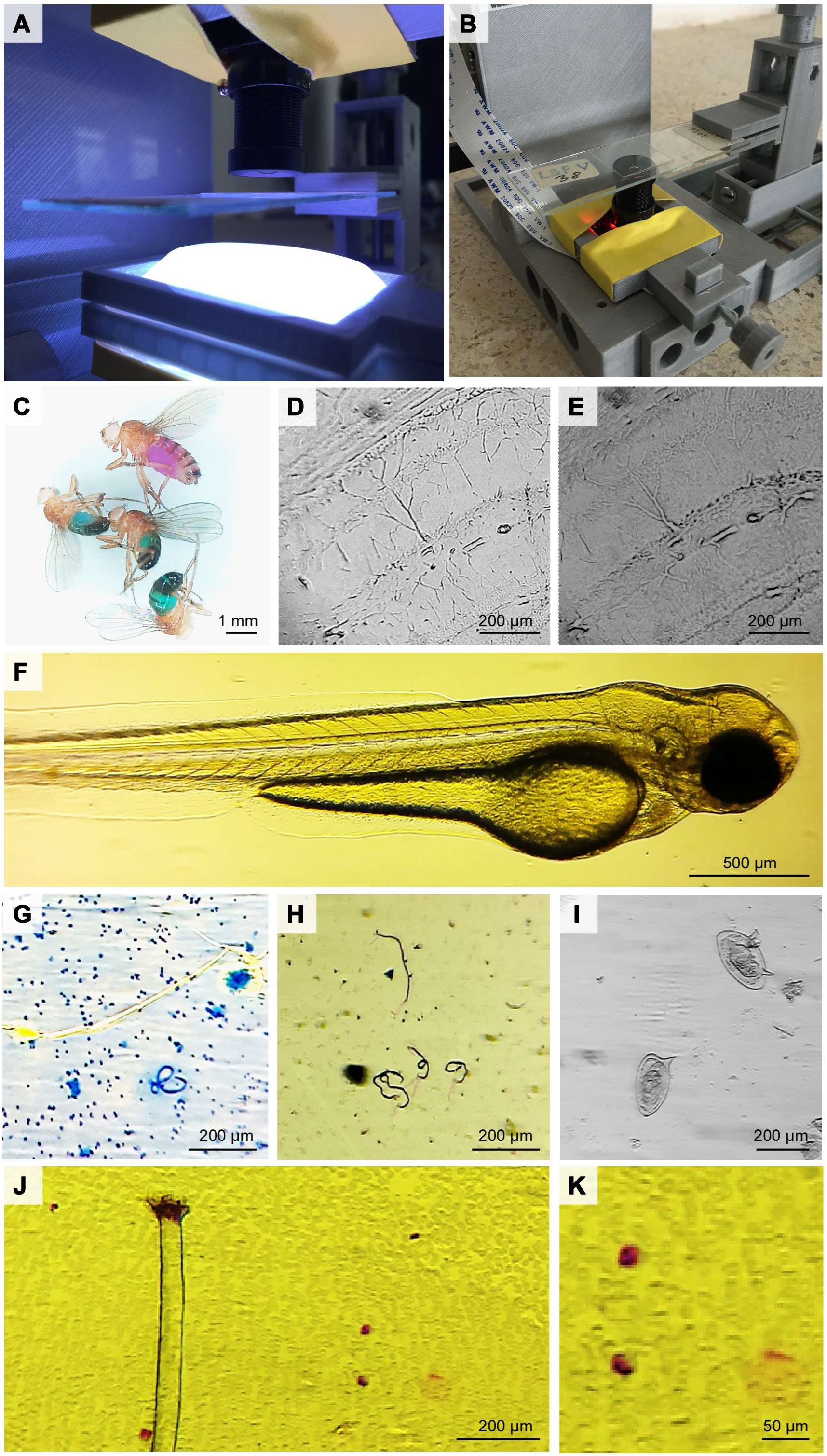
Basic Light Microscopy. **A, B**, The camera and objective can be mounted in upright (A) or inverted mode (B). In each case, the micromanipulator allows accurate positioning of a microscope slide in the image plane, while the LED ring coupled to a series of diffusors provides for flexible spectrum and brightness illumination (A). **C**, At low zoom, the magnification is appropriate to provide high-resolution colour images of several animals at once (here: *D. melanogaster* fed with fed with 5 mM sucrose in 0.5% agarose dyed with blue or red food dyes (Food Blue No. 1 and Food Red No. 106 dyes; Tokyo Chemical Industry Co., Japan) as described in [51]. **D**, **E**, When the objective is fully extended, magnification is sufficient to resolve large neurons of the mouse brain, while different positions of the LED ring permit to highlight different structures in the tissue. **F,** The system is also appropriate to provide high-resolution imagery of zebrafish larvae (D. rerio) with only room-lighting (cf. Supplementary Video 1). **G, H**, Brugia malayi (G) and Wuchereria bankrofti (H) in human lymph tissue biopsy. **I**, Schistosoma eggs in human urine. J, Mansonella perstans in human blood smear (Wright Giemsa stain) and **K,** magnification of bottom right image section.

A custom written Graphical User Interface (GUI, Supplementary Fig. 1) using the Python based PiCamera library allows for control of framerates, sensitivity, contrast, white balance and digital zoom (see Assembly and User manual in Supplementary Materials). Control over other parameters can be added as required. The GUI facilitates saving images and image sequences in jpeg format and video data in h264 or audio video interleave (AVI) format. Notably, the GUI can also function independent of the remainder of FlyPi components if only easy control for a RPi camera is required.

The camera can be mounted in two main configurations: upright or inverted (Fig. 2A, B). While the former may be primarily used for resolving larger objects such as adult *Drosophila* (Fig. 2C) or for behavioural tracking (cf. Fig. 4), the latter may be preferred for higher-zoom applications (Figs. 2D,E) and fluorescence microscopy (cf. Fig. 3), or if easy access to the top of a sample is required. Here, the image quality is easily sufficient to monitor basic physiological processes such as the heartbeat or blood-flow in live zebrafish larvae (Fig 2F, Supplementary Video 1).

**Figure 3.**
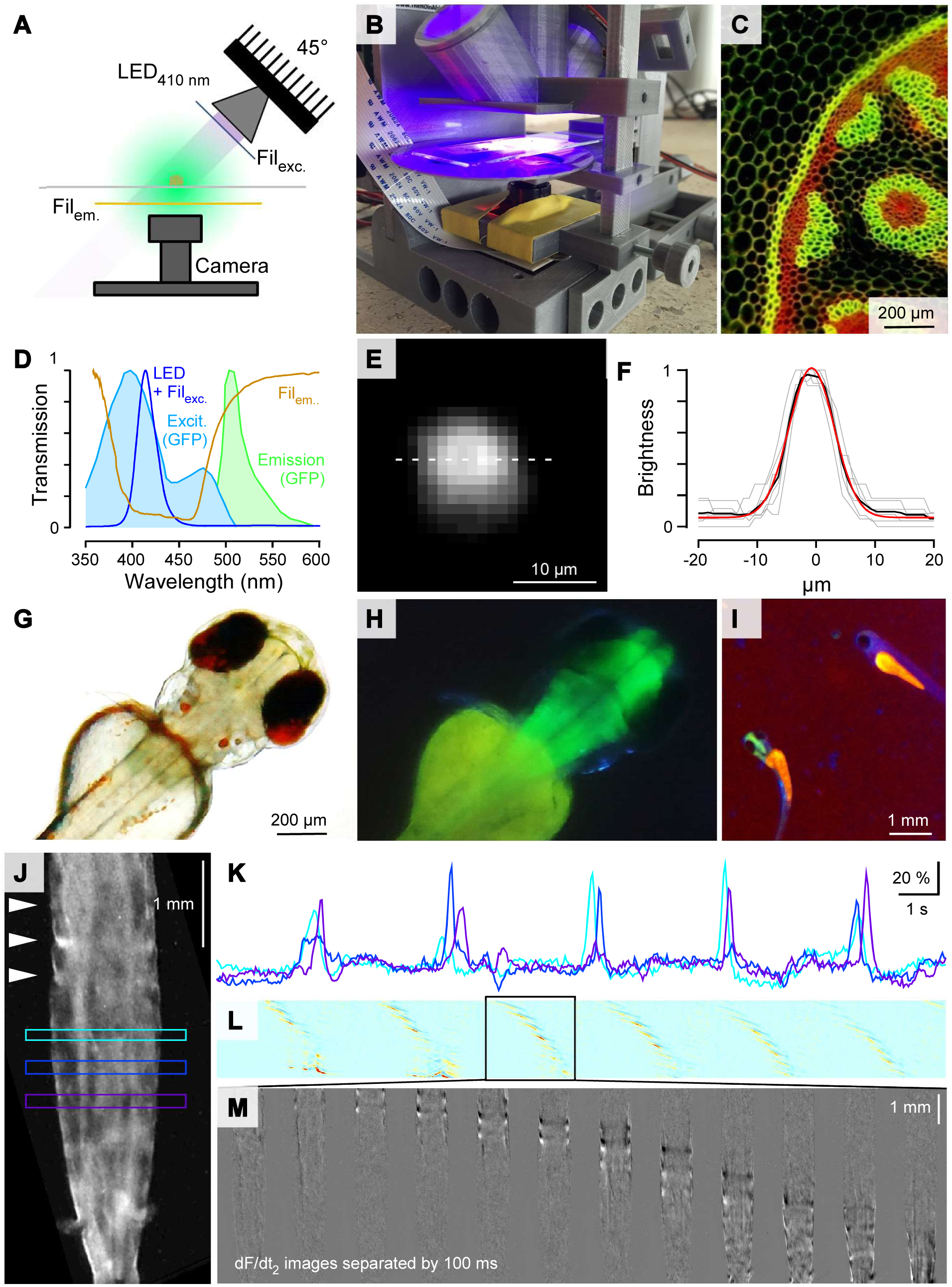
Fluorescence Microscopy. **A,** A collimated 410 nm LED angled at 45° and two ultra-low-cost theatre-lighting filters provide for fluorescence capability. **B,** Photo of the fluorescence setup. **C,** Fluorescence test-slide. **D,** Spectra of excitation LED and filters superimposed (dark blue) on GFP excitation (light blue) and emission (green) spectra. Emission filter in orange. **E, F,** Point-spread function (*psf*) measured using green fluorescent beads (Methods): Standard deviation (SD) ∼5.4 μm. **G, H,** 3 dpf Zebrafish larva expressing GCaMP5Gf in neurons (HuC:GCaMP5G) in transmission (G) and fluorescence mode (H). **I,** At low zoom the system can be used for fish-sorting (cf. Supplementary Video 2). Note the absence of green fluorescence in the brain of the non-transgenic animal to the upper right. **J-M.** Calcium Imaging in *Drosophila* larva expressing GCaMP5 in muscles (Mef2-Gal4; UAS-myr::GCaMP5). **J, K,** Three regions of interest (ROIs) placed across the raw image-stack of a freely crawling larva (J) reveal period bouts of increased fluorescence as peristaltic waves drive up calcium in muscles along the body (K). Arrowheads in J indicate positions of peaks in calcium wave. **L,** A space-time plot of the time-differentiated image stack, averaged across the short body axis, reveals regular peristaltic waves. Warm colours indicate high positive rates of change in local image brightness. **M,** A single peristaltic wave (as indicated in L) in 12 image planes separated by 100 ms intervals (cf. Supplementary Video 5).

If required, specimens can be positioned by a 3-D printed micromanipulator [4] (Fig. 2B). Up to three manipulators can be attached to the free faces of FlyPi (Fig. 1D, I). Manipulators can also be configured to hold probes such as electrodes or stimulation devices (Fig. 1I). Like the camera objective, manipulators can be optionally fitted with continuous-rotation servo motors to provide electronic control of movement in 3 axes [4]. These motors can be either software controlled, or via a stand-alone joystick-unit based on a separate Arduino-Uno microcontroller and a Sparkfun Joystick shield [17]. Depending on print quality and manipulator configuration, precision is in the order of tens of microns [18].

For lighting, we use an Adafruit Neopixel 12 LED ring [19] comprising 12 high-power RGB-LEDs that can be configured for flexible intensity and wavelength lighting. For example, the LED ring with all LEDs active simultaneously can be used to add “white” incident or transmission illumination (e.g. Fig. 2A, cf. Fig. 5B for spectra), while behavioural tracking may be performed under dim red light (cf. Fig. 4A). A series of white weighing boats mounted above the ring can be used as diffusors (Fig. 2A). Long-term time-lapse imaging, for example to monitor developmental processes or bacterial growth, can be performed in any configuration. Lighting is controlled from the GUI through an open Adafruit LED control Python library.

**Figure 4.**
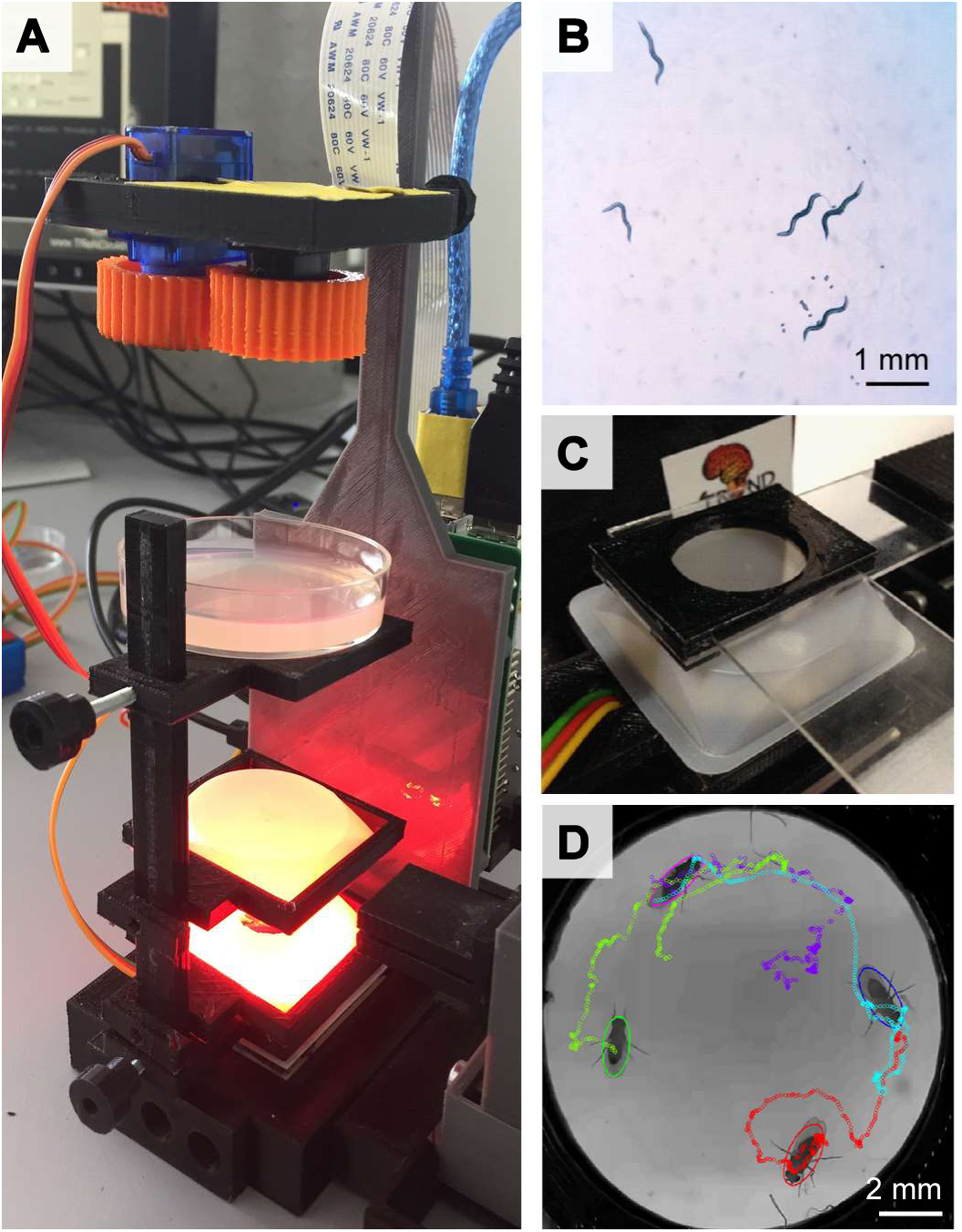
Behavioural Tracking. **A, B,** Red-light illumination from the LED ring can be used to illuminate animals during behavioural tracking – here showing *C. elegans* on an Agar plate (B). **C**, A behavioural chamber based on two microscope slides and a 3D printed chassis is adequate for behavioural monitoring of adult *Drosophila*. **D**, Animals tracked using Ctrax [23].

The implementation of a cost-effective option for digital microscopy also opens up possibilities for basic medical diagnosis, such as the detection of small parasitic nematodes *Brugia malayi* or *Wuchereria bankrofti* in human lymph tissue samples (Fig. 2G, H) or *Schistosoma* eggs in human urine (Fig. 2I). Similarly, the image is sufficient to detect and identify counterstained types of blood cells in an infected smear (here: *Mansonella perstans*; Fig. 2J, K).

### Fluorescence microscopy

Next, we implemented fluorescence capability based on a 350 mA 410 nm LED attached to a reflective collimator as well as ultra-low cost theatre-lighting filters. For this, the excitation and emission light was limited by a low-pass and a notch filter, respectively (Fig. 3A, D, Supplementary Table 1). Imperfect emission filter efficiency for blocking direct excitation light necessitated that the source was positioned at 45° relative to the objective plane, thereby preventing direct excitation bleed-through into the camera path (Figs. 3A, B). Many commonly used fluorescent proteins and synthetic probes exhibit multiple excitation peaks. For example, Green Fluorescent Protein (GFP) is traditionally excited around 488 nm, however there is a second and larger excitation peak in the near UV [20] (Fig. 3D, but see e.g. [1]). Here, we made use of this short-wavelength peak by stimulating at 410 nm to improve spectral separation of excitation and emission light despite the suboptimal emission filter. Figure 3C shows the fluorescence image recorded in a typical fluorescence test-slide. The RGB camera chip allowed simultaneous visualisation of both green and red emission. If required, the red channel could be limited either through image processing, or by addition of an appropriate short-pass emission filter positioned above the camera. Next, using green fluorescent beads (100 nm, Methods) we measured the point spread function (*psf*) of the objective as 5.4 μm (SD) at full zoom (Fig. 3E, F). This is approximately ten times broader than that of a typical state-of-the art confocal or 2-photon system [21], though without optical sectioning, and imposes a theoretical resolution limit in the order of ∼10 μm. Notably, with an effective pixel size of ∼1 μm (Fig. 1E) the system is therefore limited by the objective optics rather than the resolution of the camera chip such that the use of a higher numerical aperture objective would yield a substantial improvement in spatial resolution. It also means that at peak zoom, the camera image can be binned at x4 for increased speed and sensitivity without substantial loss in image quality.

Next, we tested FlyPi’s performance during fluorescence imaging on live animals. At lower magnification, image quality was sufficient for basic fluorescence detection as required for example for fluorescence based sorting of transgenic animals (screening). We illustrate this using a transgenic zebrafish larva (3 dpf) expressing the GFP-based calcium sensor GCaMP5G in all neurons (Fig. 3 G-I, Supplementary Video 2). Similarly, the system also provided sufficient signal-to-noise for basic calcium imaging, here demonstrated using *Drosophila* larvae driving GCaMP5 in muscles that reveal clear fluorescence signals associated with peristaltic waves as the animal freely crawls on a microscope slide (Fig. 3J-M; see also Supplementary Video 3, cf. [22]). Further fluorescence example videos are provided in the supplementary materials (Supplementary Videos 4,5).

### Behavioural tracking

“To move is all mankind can do”. Sherrington’s (1924) thoughts on the ultimate role of any animal’s nervous system still echoes today, where despite decades of (bio)technological advances, behavioural experiments are still amongst the most powerful means for understanding neuronal function and organisation. Typically, individual or groups of animals are placed in a controlled environment and filmed using a camera system. Here, FlyPi’s colour camera with adjustable zoom offers a wide range of video-monitoring options, while the RGB LED ring provides for easily adjusted wavelength and intensity lighting (Fig. 4A) including dim red light, which is largely invisible to many invertebrates including *C. elegans* (Fig. 4B, Supplementary Video 6) and *Drosophila.* A series of mounting adapters for petri-dishes (Fig. 1H) as well as a custom chamber consisting of a 3-D printed chassis and two glass microscope slides for adult *Drosophila* (Fig. 4C) can be used as behavioural arenas. Following data acquisition, videos are typically fed through a series of tracking and annotation routines to note the spatial position, orientation or behavioural patterns of each animal. Today, a vast range of open behavioural analysis packages is available, including many that run directly on the RPi3 such as CTrax [23], here used to track the movements of adult *Drosophila* in a 10 s video (Fig. 4D; Supplementary Video 7).

### Optogenetics and Thermogenetics

One key advantage of using genetically tractable model organisms is the ability to selectively express proteins in select populations of cells whose state can be precisely controlled using external physical stimuli such as light (Optogenetic effectors, e.g. [24]) or heat (Thermogenetic effectors, e.g. [15]). Through these, the function of individual or sets of neurons can be readily studied in behavioural experiments. A plethora of both light- and heat- sensitive proteins are available, with new variants being continuously developed. Many of these proteins exhibit sufficient sensitivity for activation by collimated high-power LEDs, rather than having to rely on more expensive light sources like a Xenon lamp or a laser. Similarly, temperature variation over few degrees Celsius, as achieved by an off-the-shelf Peltier element with adequate heat dissipation, is sufficient to activate or inactivate a range of temperature-sensitive proteins. We therefore implemented both opto- and thermogenetic stimulation capability for FlyPi.

*Optogenetics.* For optogenetic activation we used the LED ring (Fig. 5A), whose spectrum and power are appropriate for use with both ChR2 (single LED ‘blue’ Pwr_460_: 14.2 mW) as well as ReaChr and CsChrimson (‘red’ Pwr_628_: 7.2 mW; ‘green’ Pwr_518_: 7.5 mW) (Fig. 5B) [3], [14], [25], [26]. Alternatively, an Adafruit 8×8 high-power single wavelength LED matrix [19] can be attached for spatially selective optogenetic or visual stimulation [27]. For demonstration, a zebrafish larva (3 dpf) expressing ChR2 in all neurons was mounted on top of a microscope slide, which was in turn held above the inverted objective using the micromanipulator (Fig. 5A, C). The LED ring was positioned face-down ∼2 cm above the animal, outside of the centrally positioned camera’s the field of view. Concurrent maximal activation of all 12 ‘blue’ LEDs (Pwr_460_: ∼4.9 mW cm^-2^ at the level of the specimen) reliably elicited basic motor patterns for stimuli exceeding 500 ms, here illustrated by pectoral fin swimming bouts (Fig. 5C,D, Supplementary Video 8). Substantially shorter stimuli did not elicit the behaviour (e.g. 3^rd^ trial: ∼150 ms), nor did activation of the other wavelength LEDs or blue light activation in ChR2-negative control animals (not shown). This strongly indicated that motor networks were activated through ChR2 rather than innate visually-mediated escape reflexes in response to the light (cf. [28]) or photomotor responses [29]. Notably, while in the example shown the stimulus artefact was used as a timing marker, excitation light could be blocked (>95% attenuation) using an appropriate filter (Fig. 5B dark red trace, Supplementary Table 1) without substantially affecting image quality, while timing could be verified using the flexibly programmable low-power RGB LED normally integrated into the Peltier-thermistor loop (not shown).

**Figure 5.**
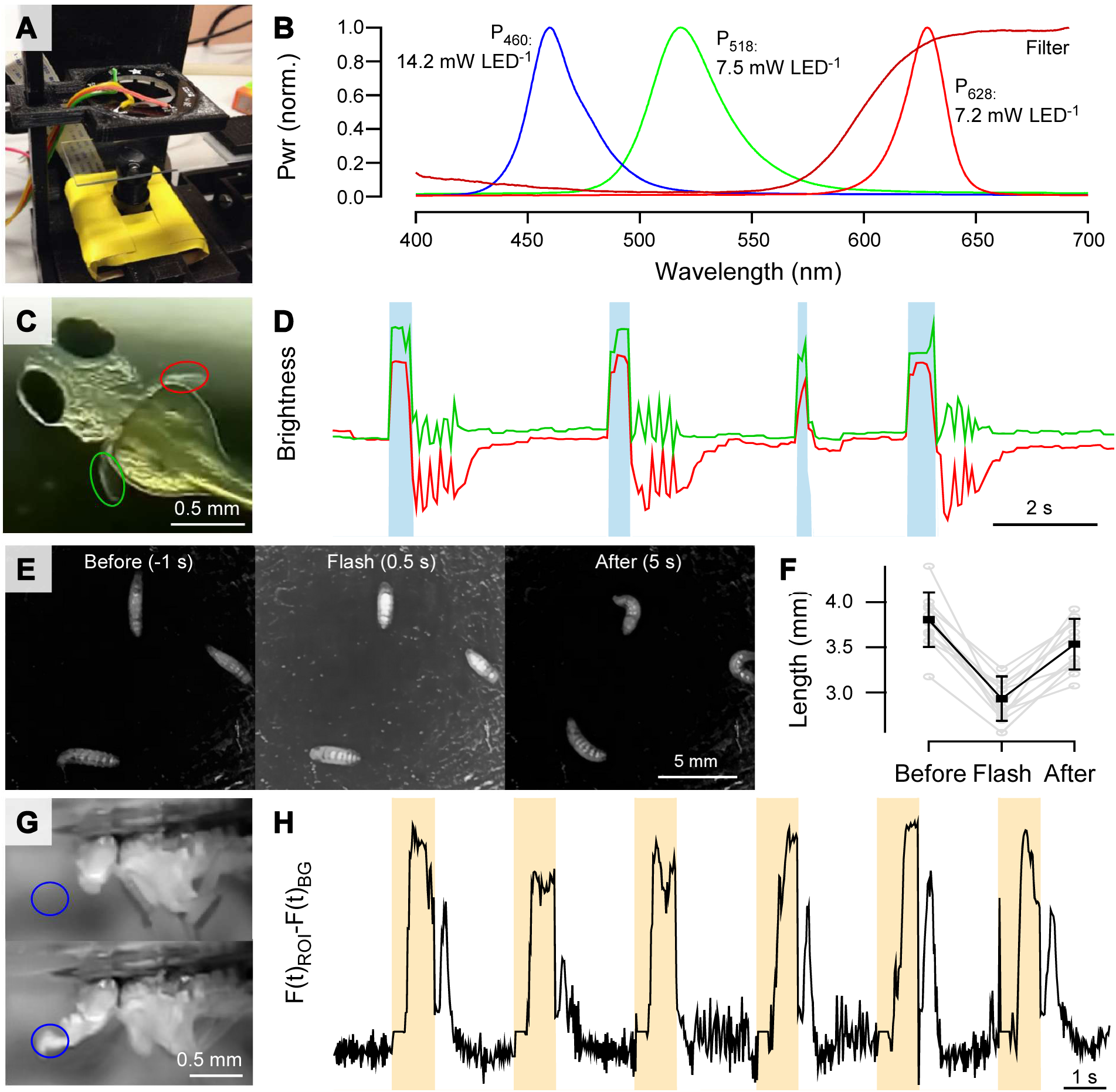
Optogenetics. **A,** Experimental configuration suitable for optogenetic stimulation of an individual zebrafish larva suspended in a drop of E3. **B**, Spectrum and peak power of the three LEDs embedded at each ring position. Spectral filters can be used to limit excitation light reaching the camera (Rosco Supergel No. 19, “Fire”). **C**, zebrafish larva (3 *dpf*) expressing ChR2 broadly in neurons (Et(E1b:Gal4)s1101t, Tg(UAS::Cr.ChR2_H134R-mCherry) s1985t, nacre-/-). **D**, The animal exhibits pectoral fin burst motor patterns upon activation of blue LEDs (cf. Supplementary Video 8). **E, F**, *Drosophila* larvae expressing ChR2 in all neurons (elav-GAL4/+; UAS-shibre^ts^; UAS-ChR2/+; UAS-ChR2/+) crawling on ink-stained agar reliably contract when blue LEDs are active. **G, H**, Proboscis extension reflex (PER) in adult *Drosophila* expressing CsChrimson in the gustatory circuit (courtesy of Olivia Schwarz and Jan Pielage, Friedrich Miescher Institute for Biomedical Research, Basel, *unpublished line*) is reliably elicited by activation of red LEDs.

We also tested ChR2 activation in *Drosophila* larvae. Animals were left to freely crawl on ink-stained agarose with both the LED ring and camera positioned above. Activation of all 12 blue LEDs reliably triggered body contractions for the duration of the 1 s stimulus, followed by rapid recovery (Fig. 5E, F). Finally, full-power activation of the red LEDs reliably triggered proboscis extension reflex (PER) in adult *Drosophila* expressing CsChrimson in the gustatory circuit (Fig. 5G,H). In this latter demonstration, we made use of the GUI’s protocol function which allows easy programming of microsecond-precision looping patterns controlling key FlyPi functions such as LEDs and the Peltier Loop, (cf. Fig. 6C).

**Figure 6.**
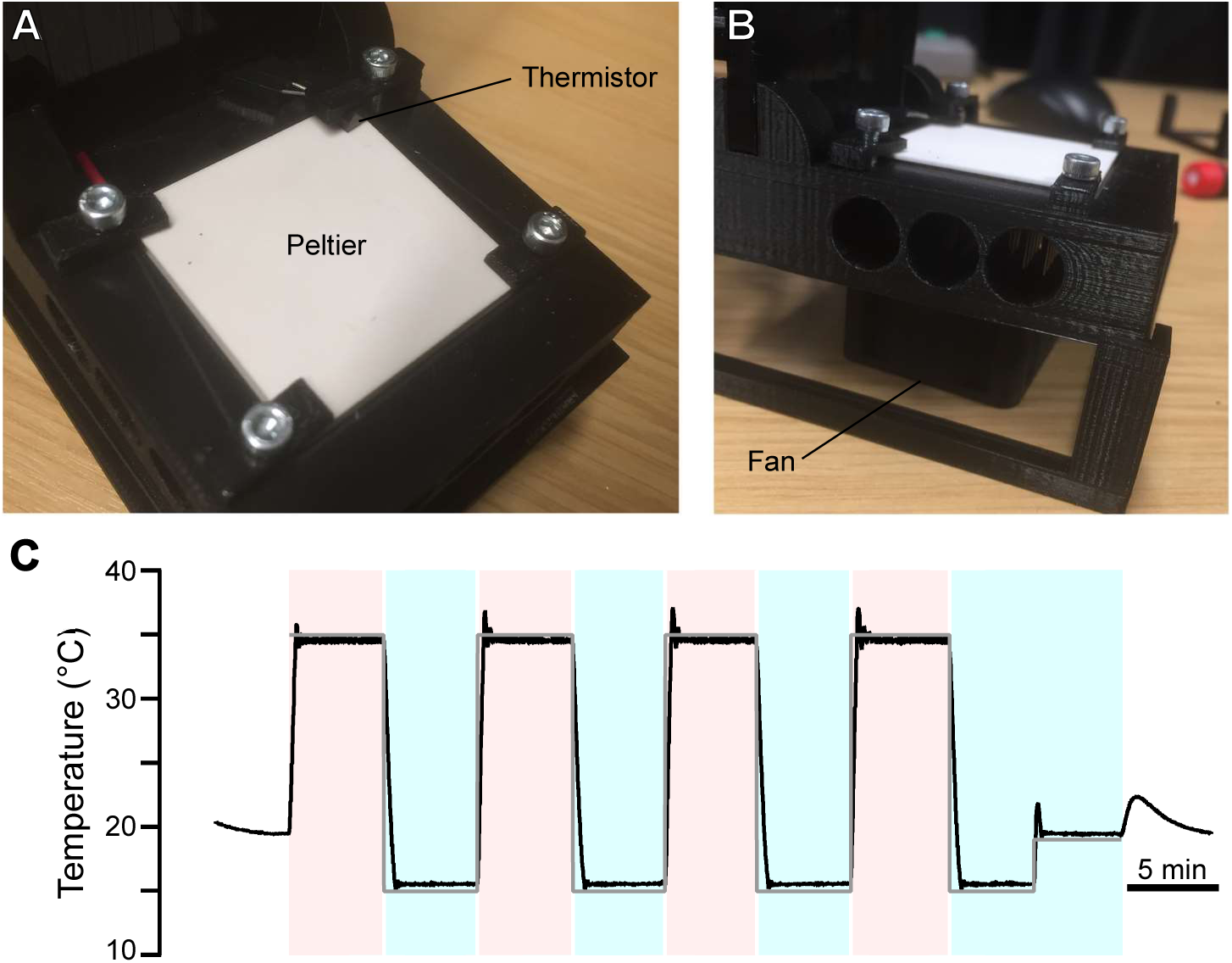
Thermogenetics. **A**, The 4×4 Peltier element embedded in the FlyPi base, with the Thermistor clamped into one corner. **B**, Side-view with FlyPi propped up on a set of 3D printed feet to allow air dissipation beneath the base. The CPU fan is positioned directly beneath the Peltier. **C**, Performance of the Peltier-thermistor feedback loop. Command 15°C and 35°C indicated by blue and red shading switching every 5 mins; room temperature 19°C (no shading).

*Thermogenetics.* Owing to their remarkable ability to tolerate a wide range of ambient temperatures, many invertebrate model species including *Drosophila* and *C. elegans* also lend themselves to thermogenetic manipulation. Through the select expression of proteins such as Trp-A or shibire^ts^ [15], [30], sets of neurons can be readily activated or have their synaptic drive blocked by raising the ambient temperature over a narrow threshold of 28 and 32°C, respectively. Here, FlyPi offers the possibility to accurately control temperature of the upper surface of a 4×4cm Peltier element embedded in its base, with immediate feedback from a temperature sensor (Fig. 6A, Supplementary Table 1). A CPU fan and heat sink below the Peltier element dissipate excess heat (Fig. 6B, Supplementary Table 1). The setup reaches surface temperatures +/− ∼20°C around ambient temperature within seconds (∼1°C/s) and holds set temperatures steady over many minutes (SD <1°C) (Fig. 6C).

## DISCUSSION

We primarily designed FlyPi to achieve a good balance of performance and cost and flexibility in its use. Using higher quality components, individual function performance can certainly be improved (see *Potential for further development*). Here, it is instructive to compare FlyPi’s microscope function to other open microscope designs. For example, the fully 3-D printable microscope stage of the “Waterscope” [31] achieves superior stability of the focussing mechanism. However, unlike FlyPi, this design cannot achieve the same range of possible magnifications needed for behavioural experiments. Some other open microscope designs (e.g. [32]) use a larger fraction of commercial components of provide superior image quality and/or stability, albeit invariably at substantially higher cost. On the extreme low-cost scale, available designs typically do not provide the imaging systems itself (i.e. the camera, control software and processor) but instead rely on the addition of a mobile phone camera or, indeed, the eye itself (e.g.[33], [34]). Next, FlyPi also provides for a powerful range of sample illumination options, which typically exceed available alternatives. Importantly, to our knowledge, no alternative open-microscope design encompasses the experimental accessories and control systems required for behavioural tracking under neurogenetic control.

Another key aspect of FlyPi’s design is its modular nature. This means that the system does not require all integrated options to be assembled to function. For example, if the main purpose of an assembled unit is to excite Channelrhodopsin, the only module beside the base unit is the LED ring. Similarly, only the Peltier-thermistor circuit is needed for Thermogenetics experiments. This means that units designed for a dedicated purpose can be assembled quickly and at substantially reduced cost. Moreover, given a functional base unit, it is easy for the user to modify any one part or to integrate a fully independent module built for a different purpose altogether. The modular nature also renders the design more robust in the face of difficulties with sourcing building components.

### Potential for further development

Clearly, the current FlyPi only scratches the surface of possible applications. Further development is expected to take place as researchers and educators integrate aspects of our design into their laboratory routines. To explicitly encourage re-sharing of such designs with the community we maintain and curate a centralised official project page (http://open-labware.net/projects/flypi/) linked to a code repository (https://github.com/amchagas/Flypi). Indeed, a basic description of the FlyPi project has been online since 2015 which has led to several community-driven modifications. For example, a recent modification of the 3D printed mainframe implements the camera and focus motor below a closed stage [35]. At the expense of a fixed camera position, this build is substantially more robust and thus perhaps more suitable e.g. for classroom teaching. Other community driven modifications include a version where all 3D printed parts are replaced by Lego^®^ blocks [36] as well as several forks geared to optimize the code, details in the 3D model or additions in the electronic control circuits.

Currently, one obvious limit of FlyPi is spatial resolution. The system currently resolves individual human red blood cells (Fig. 2K), but narrowly fails to resolve malaria parasites within (not shown). Here, the limit is optical rather than related to the camera chip, meaning that use of a higher numerical aperture and magnification objective lens will yield substantial improvements. This development might come in hand with additional improvements in the micromanipulator’s Z-axis stability to facilitate focussing at higher magnification - for example as implemented in the Waterscope [31]. Similarly, photon catch efficiency of the CCD sensor could be improved by use of an unfiltered (monochrome) chip. Other alleys of potential further development include (i) the addition of further options for fluorescence microscopy to work over a wider range of wavelengths, likely through use of other excitation LEDs and spectral filters. (ii) FlyPi could also be tested for stimulating photo-conversion of genetically encoded proteins such as CamPari, Kaede or photoconvertible GFP [37]–[39]. (iii) Auto-focussing could be implemented by iteratively rotating the servo-assisted focus while evaluating changes in the spatial autocorrelation function or Fourier spectrum of the live image. (iv) A motorised manipulator could be integrated for stage-automation through a simple software routine. (v) One or several FlyPis could be networked wirelessly or through the integrated Ethernet port to allow centralised access and control, thereby removing need for dedicated user interface peripherals. Taken together, by providing all source code and designs under an open source license, together with an expandable online repository, we aim to provide a flexible, modular platform upon which enthusiastic colleagues may build and exchange modifications in time.

### Classroom teaching and laboratory improvisation

In large parts of the world, funding restrictions hamper the widespread implementation of practical science education – a problem that is pervasive across both schools and universities [18], [40]. Often, limitations include broken or complete lack of basic equipment such as low power light microscopes or computing resources. Here, the low cost and robustness of FlyPi may offer a viable solution. If only one unit can be made available for an entire classroom, the teacher can project the display output of FlyPi to the wall such that many students can follow demonstrated experiments. Already a low amount of funding may furnish an entire classroom with FlyPis, allowing students in pairs of two or three to work and maintain on their own unit. The relative ease of assembly also means that building FlyPi itself could be integrated into part of the syllabus. In this way, a basic technical education in electronics and soldering or basic 3-D printing could be conveyed in parallel. As an additional advantage, each student could build their own equipment which brings about further benefits in equipment maintenance and long-term use beyond the classroom.

To survey to what extent FlyPi assembly and use may be beneficial in a classroom scenario, we introduced the equipment to African biomedical MSc and PhD students as well as senior members of faculty during a series of multi-day workshops at Universities in sub-Saharan Africa since 2015, including the University of KwaZulu Natal (Durban, SA), the International Centre of Insect Physiology and Ecology (icipe, Nairobi, Kenya), Kampala International University (Dar es Salaam, Tanzania and Ishaka Bushyeni, Uganda) and the International Medical and Technical University (IMTU, Dar es Salaam, Tanzania). In addition, colleagues have used and modified the design for projects held in Accra, Ghana, Khartoum, Sudan and Ishaka, Uganda. In one workshop, we only provided the 3-D printed parts, the custom PCB and off-the-shelf electronics and took students though the entire process of assembly and installation (Fig. 7A). Having had no previous experience with basic electronics, soldering or the use of simple hand-tools such as a Dremel or cable-strippers, all students successfully assembled a working unit. Towards the end of the training, students used their own FlyPi to perform basic Neurogenetics experiments with *Drosophila*, including heat activation of larvae expressing shibire^ts^ in all neurons (elav-GAL4/+; UAS-shibre^ts^,UAS-ChR2 / +; UAS-ChR2 / +, cf. Fig. 6) and optogenetic activation of ChR2 to elicit a range of behaviours in both adults and larvae (cf. Fig. 5). Following the training, students took their assembled FlyPis home for their own research and teaching purposes. In other courses, we brought pre-assembled FlyPis with a range of different modules. Students learnt to operate the equipment within minutes and subsequently used them for a range of experiments and microscopy tasks, including several novel configurations not formally introduced by the faculty (Figs. 7B, C). Indeed, many experiments presented in this paper were performed during these training courses. Finally, we used individual FlyPi modules to improvise workarounds for incomplete commercial lab equipment. For example, the RPi camera with focus drive and live image-processing options served as an excellent replacement for a missing Gel-doc camera (Fig. 7D). Similarly, we used FlyPi as a replacement camera for odour evoked calcium imaging in *Drosophila* antennas on a commercial upright fluorescence microscope or for dissection demonstrations under a stereoscope that also utilised the LED rings for illumination. Moreover, FlyPi’s programmable General Purpose Pins (GPPs) and LEDs were used to drive time-precise light-stimulus series, e.g. for independently recorded *Drosophila* electroretinograms (ERGs). Similarly, the Peltier-feedback circuit was adequate to maintain developing zebrafish embryos at a controlled temperature during prolonged experiments, or to reversibly block action potential propagation in long nerves through local cooling. Clearly, beyond its use as a self-standing piece of equipment and teaching tool, the low cost and modular nature of FlyPi also renders it versatile to support or take over a large range of additional functions in the lab.

**Figure 7.**
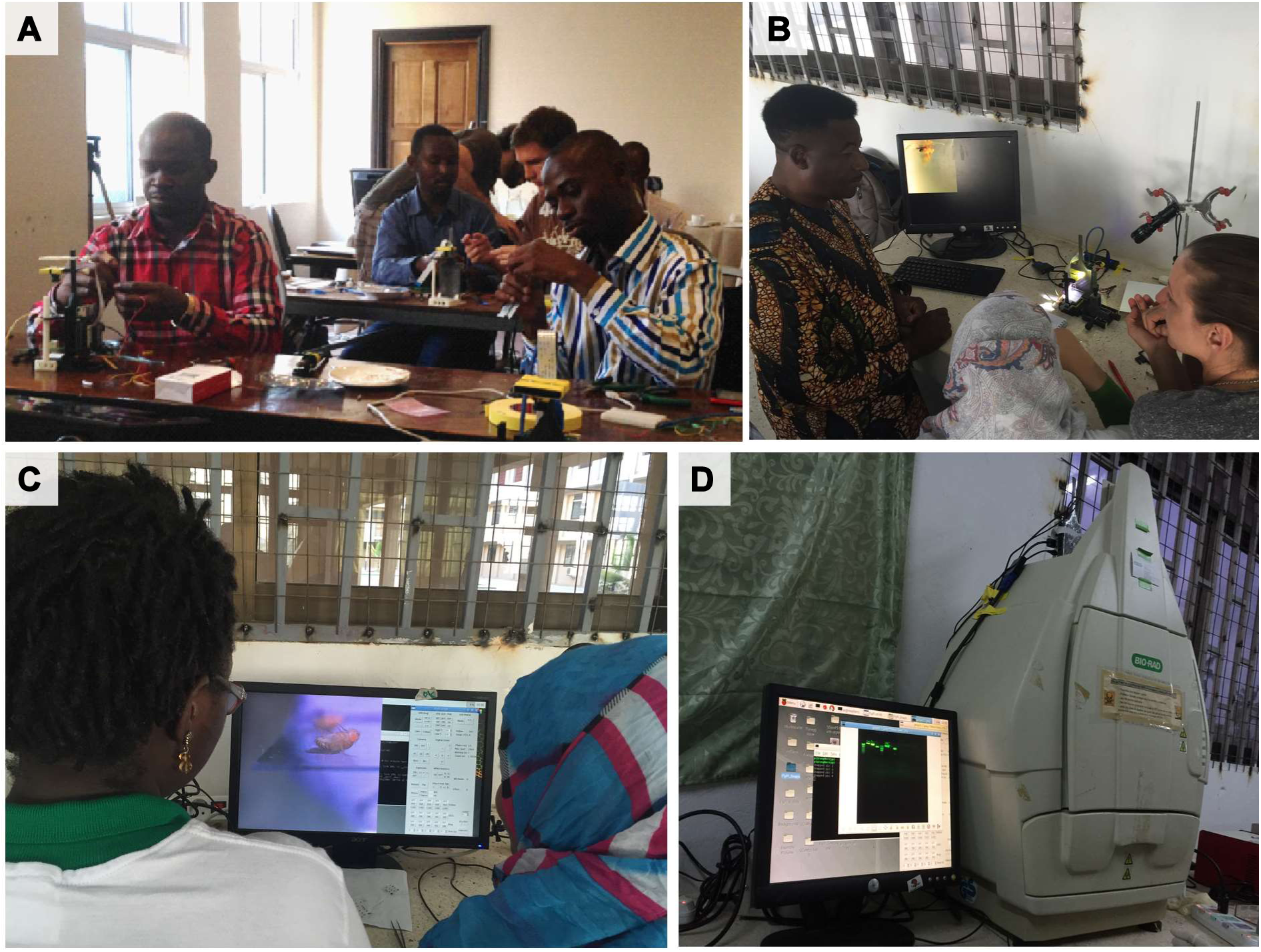
Classroom teaching and equipment improvisation. **A**, Graduate students from different African Universities building FlyPis during a workshop held in Durban, South Africa in March 2015. **B, C** African graduate students and faculty in Dar es Salaam, Tanzania, using FlyPis for optogenetics experiments on proboscis extension reflex as readout. **D**, FlyPi with adjustable focus module mounted on top of a Gel-Doc used to replace missing commercial camera.

## CONCLUSION

Taken together, we anticipate that the open design of FlyPi will be useful in scientific teaching and research as well as for medical professionals working in low-resource settings looking to supplement their diagnostic toolkit. We anticipate that in time, further improvement and new designs will emerge from the global open hardware community. Notably, a curated collection of further such “Open-Labware” [18], [41] designs can be found on the PLoS website [42].

## METHODS

### A complete assembly and user manual is provided in the Supplementary Materials

#### 3D Modelling and printing

3-D modelling was performed in OpenSCAD [43] and all files are provided as both editable scad and complied surface tesselation lattice (stl) files. All parts were printed in polylactic acid (PLA) on an Ultimaker 2 3-D printer (Ultimaker, Geldermailsen, Netherlands) in six pre-arranged plates using the following parameters: infill 30%, no supports, 5 mm brim, layer height 0.1 mm, print speed 60 mm/s, travel speed: 200 mm/s. Total printing time of a single FlyPi, including all presented modules, was about 40 hours. Notably, this time can be substantially reduced by using faster print settings and/or a larger nozzle, as e.g. commonly implemented in lower-cost 3-D printers. For example, using a well-calibrated delta Rep-Rap delta (www.reprap.org) printing at full speed, the entire system can be printed at sufficient precision in less than 20 hours.

#### PCB design and printing

The printed circuit board (PCB) was designed in KiCad [44] and is provided as the native KiCad file format as well as the more widely used gerber file format. The PCB facilitates connections between peripherals and the microcontroller, and was designed to be modular such that only components that will be used need to be soldered on the board. The power circuitry designed for one single 12 V 5 A power supply is provided. The large spacing between component slots, PCB labelling and consistent use of the “through-hole” component format is intended to facilitate assembly by users with little soldering experience. Using the provided Gerber files, it is possible to order the PCBs from a variety of producers (e.g., pcbway.com, seeedstudio.com/pcb, dirtypcbs.com). Of course, if required the entire PCB could also be improvised using individual cables and/or a suitable breadboard by taking reference to the circuit diagram provided.

#### The Graphical User Interface (GUI)

The GUI (Supplementary Fig. 1) was written in Python3. The control functions for each peripheral component is created in its own class, making it easier for the end user to create/alter functions independently. These classes are then contained in a “general purpose” class, responsible for the display of the user interface and addressing the commands to be send to the Arduino board (responsible for time-precise events and direct interaction with peripherals, for details see below). The communication between the RPi and the Arduino is established via universal serial bus (USB) through a serial protocol (Python Serial library [45]). The GUI is created using Tkinter [46]. Both libraries are compatible with Python2 and Python3.

The GUI is also capable of creating folders and saving files to the Raspberry Pi desktop. For simplicity, the software creates a folder called “FlyPi_output” and subfolders depending on the type of data being acquired (time lapse, video, snapshots, temperature logging). The files within the subfolders are created using date and time as their names, preventing overwriting of data.

#### Arduino

We used an ATmega328 based Arduino Nano [12]. The board was chosen due to a high number of input/output ports, its variety of communication protocols (e.g. Serial, I2C), its low cost and easy availability (including several ultra-low cost clones at 2-3 €), very well documented environment (hardware specifications, function descriptions, “how to” recipes), and large user database. The board is programmed in C++ together with the modifications added by the Arduino integrated development environment (IDE). The board is responsible for controlling all peripheral devices except the camera, and provides microsecond precision for time measurement. The code can be adapted to most of the other boards of the Arduino family, with small changes (e.g., digital, analogue and serial port addresses).

*Raspberry Pi 3 operating system.* We used Raspian [47] as the operating system (OS) on the Raspberry Pi 3 [13] for its installation simplicity through “new out of the box software” (NOOBS) [48] and because it is derived from Debian [49], a stable and well-supported GNU-Linux distribution. However, any Linux distribution compatible with the Raspberry Pi and the chosen Python3 libraries can be used. Arduino compatibility is not mandatory, since once the board is loaded with the correct code, which can be done in any computer, the Arduino IDE is not used further as all live communication goes via the serial port directly from Python.

#### Spectral and power measurements

We used a commercial photo-spectrometer (USB2000+VIS-NIR, Ocean Optics, Ostfildern, Germany) and custom written software in Igor-Pro 7 (Wavemetrics) to record and analyse spectra of LEDs and filters. Peak LED power was determined using a Powermeter (Model 818, 200-1800 nm, Newport). We used fluorescent beads (PS-Speck TM Microscope Point Source Kit P-7220, ThermoFisher) for estimating FlyPi’s point spread function (*psf*).

#### Video and image acquisition

All static image data was obtained as full resolution red-green-blue (RGB) images (2592×1944 pixels) and saved as jpeg. All video data was obtained as RGB at 42 Hz (x2 binning), yielding image stack of 1296×972 pixels, and saved as h264. Video data was converted to AVI using the ffmpeg package for GNU/Linux (ffmpeg.org, a conversion button is added to the GUI for simplicity). All further data analysis was performed in Image-J (NIH) and Igor-Pro 7 (Wavemetrics). Figures were prepared in Canvas 15 (ACD Systems).

#### Calcium imaging in larval Drosophila muscles

Second instar larvae (Mef2-Gal4; UAS-myr::GCaMP5) were left to freely crawl between a microscope slide and cover slip loosely suspended with tap water. For analysis, x2 binned video data (42 Hz) was further down-sampled by a factor of 2 in the image plane and a factor or 4 in time. Only the green channel was analysed. Following background subtraction, regions of interest were placed as indicated (Fig. 3J). Next, from each image frame we subtracted the mean image of 4 preceding frames to generate a “running average time-differential” stack – shown as the space-time plot in Fig. 3L with the original x-axis collapsed. Individual non-collapsed frames of this stack, separated by 100 ms intervals, are shown in Fig. 3M.

#### Zebrafish ChR2 activation

A 3 *dpf* zebrafish larva (*Et*(*E1b:Gal4*)*s1101t*, *Tg*(*UAS:Cr.ChR2_H134R-mCherry*)*s1985t*, *nacre-/-*) was mounted in a drop of E3 medium (5 mM NaCl, 0.17 mM KCl, 0.33 mM CaCl2 0.33 mM MgSO_4,_ pH adjusted to 7.4 using NaHCO_3_) on top of a microscope slide and placed immediately above the inverted camera objective. The NeoPixel 12 LED ring was placed about 2 cm above the specimen, facing down. Concurrent maximal activation of all 12 blue LEDs for more than 500 ms reliably elicited pectoral fin swimming bouts. Shorter stimuli were not effective. RGB image data was obtained at 42 Hz, down-sampled by a factor of 4 in time and visualised by tracking the mean brightness of two regions of interest placed onto the pectoral fins.

#### Drosophila larva ChR2 activation

1^st^ instar *Drosophila* larvae (elav-GAL4/+; UAS-shibre^ts^; UAS-ChR2/+; UAS-ChR2/+) were placed on agarose darkened with Indian ink (1% v/v) within the lid of a 50 ml falcon tube and left to freely crawl. The camera and NeoPixel LED ring was placed about 3 cm above the surface. Concurrent activation of all 12 blue LEDs for 1 s at a time reliably triggered larval contractions. Image data acquired at 42 Hz and saved as 8-bit greyscale. Larval length was quantified manually in ImageJ by measuring the distance between head and tail along the body axis at 3 time-points: t = −1, 0.5 and 5 s relative to the flash (t = 0-1 s). n = 12 responses from 3 animals, error bars in SD.

#### Drosophila adult Chrimson activation

Adult *Drosophila* (Pielage, *unpublished line:* Gal4/+;UAS-CsChrimson/+, raised on standard food mixed with 200 μM all-trans retinal as described in [50]) were fixed to a cover slide by gluing the back of their thorax nail varnish, with limbs moving freely. The NeoPixel 12 LED ring was positioned around the camera objective about 2 cm above the fly pointing down. Concurrent maximal activation of all 12 red LEDs for 1 s, separated by 2 s intervals, reliably elicited the proboscis extension reflex. RGB image data was obtained at 42 Hz (x2 binning). The image stack was converted to 8-bit greyscale and background over time was subtracted from the entire image stack to limit the excitation light artefact. To calculate proboscis position over time, we plot image brightness over time within a region of interest placed at the tip of the fully extended proboscis.

#### Thermogenetics

To assess the performance and stability of the Peltier-Thermistor loop we exported the Peltier command setting and Thermistor reading at 2 Hz through the serial port into an Ascii file and analysed the data using IgorPro 6 (Wavemetrics).

## ACKNOWLEDGEMENTS

We thank the following who kindly provided technical advice, specimens, reagents and other vital forms of support: Thomas Euler (technical advice and support for A.M.C.), Ihab Riad (discussions, development and field testing), Paul Szyska (advice on optical filters), Cornelius Schwartz (support for A.M.C.), Thirumalaisamy P Velavan (human parasite slides), Della David (C. *elegans*), Marta Rivera-Alba (CTrax and field testing), Olivia Schwartz and Jan Pielage (Gal4 and UAS-CsChrimson flies and field testing), Matthias Landgraf (Mef2-Gal4; UAS-myr::GCaMP5 flies and field testing), Richard Benton (Several *Drosophila* lines, support for L.P.G.), Juan Sanchez (*Drosophila* feeding assay and field testing), Miroslav Róman-Róson (brain slice) and Georg Raiser, Tom Laudes, Lukas v Tobel and Christen Mirth (field testing). This work was supported by the Deutsche Forschungsgemeinschaft (BA 5283/1-1 to T.B), the BW-Stiftung (AZ 1.16101.09 to T.B.), the intramural fortüne program of the University of Tübingen (2125-0-0 to T.B.) and the European Commission (H2020 ERC-StG 677687 ‘NeuroVisEco’ to T.B.). A.B.A was supported by EXC307 (CIN-Werner Reichardt Centre for Integrative Neuroscience). A.M.C. was supported by the National Institute of Neurological Disorders and Stroke (U01NS090562) of the National Institutes of Health. In addition, we thank the many students and teaching volunteers as well as funders of TReND in Africa’s (www.TReNDinAfrica.org) workshops and other activities who jointly supported the inspiration for and development of FlyPi: International Brain Research Organisation, The Company of Biologists, The Wellcome Trust, The VolkswagenStiftung, The International Society of Neurochemistry, The Cambridge Alborada Trust, Cambridge in Africa, The American Physiological Society, The Physiological Society, The University of Lausanne Officine de egalite, and many others. The content is solely the responsibility of the authors and does not necessarily represent the official views of the funders.

## AUTHOR CONTRIBUTION

FlyPi was jointly designed and implemented by A.M.C and T.B with help from L.P.G. All Python and Arduino code, as well as all electronics were implemented by A.M.C with help from T.B. The OpenSCAD model was written by T.B. with help from A.M.C. T.B performed experiments and analysis, with help from all authors. The paper was written by T.B, A.M.C, L.P.G and A.B.A.

## SUPPLEMENTARY FIGURE 1 – Graphical User Interface (GUI)

Screenshots of the Python-based GUI divided into four main control panels that can be individually activated depending on user requirements: **A**, Camera control, **B**, LED, **C**, Peltier and Focus Servo control, **D**, Custom protocol window. For details, please consult the Supplementary Assembly and User Manual.

## SUPPLEMENTARY TABLE 1 – Bill of Materials (BOM)

Complete list, estimated costs and online links to all required parts, organised by modules. For details, please consult the Supplementary Assembly and User Manual.

## SUPPLEMENTARY ASSEMBLY AND USER MANUAL

Complete Assembly and User Manual

## SUPPLEMENTARY VIDEOS

1. Zebrafish larva transmission to visualise circulation (related to Fig. 2F)
2. Zebrafish larva fluorescence sorting (related to Fig. 3I)
3. Zebrafish larva expressing GFP in the heart (related to Fig. 3)
4. Zebrafish eggs expressing GCaMP5 in all neurons (related to Fig. 3)
5. *Drosophila* larva calcium imaging (related to Fig. 3J-M)
6. *C. elegans* crawling freely (related to Fig. 4B)
7. *Drosophila* adults walking freely in custom chamber (related to Fig. 4D)
8. Zebrafish expressing ChR2 in all neurons under blue light (related to Fig. 5C,D)
9. *Drosophila* larvae ChR2 under blue light (related to Fig. 5E,F)
10. *Drosophila* adult proboscis extension reflex driven by CsChrimson using red light (related to Fig. 5G,H)

